# Fine-grained neural coding of bodies and body parts in human visual cortex

**DOI:** 10.1101/2024.02.09.579107

**Authors:** Jesus Garcia Ramirez, Michael Vanhoyland, Ratan N. Apurva Murty, Thomas Decramer, Wim Van Paesschen, Stefania Bracci, Hans Op de Beeck, Nancy Kanwisher, Peter Janssen, Tom Theys

## Abstract

The visual image of a human body provides a valuable source of socially relevant information. However, our understanding of the neuronal mechanisms underlying body perception in humans remains limited given the spatiotemporal constraints of functional imaging. Here we recorded multi-unit spiking activity in two neurosurgical patients in or near the extrastriate body area (EBA), a critical region for body perception. Our recordings revealed a strong preference for human bodies over a large range of control stimuli. Notably, this preference was driven by a distinct selectivity for body parts. Moreover, the observed body selectivity generalized to non-photographic depictions of bodies such as silhouettes and stick figures. Overall, our study provides an unprecedented access into the representation of bodies in the human visual cortex to bridge the gap between human neuroimaging and macaque electrophysiology studies, and form a solid basis for computational models of human body processing.

## Introduction

Perceiving bodies is a fundamental component of social vision in humans and animals, and accordingly engages a large region of cortex specialized for this process^1^. However, the actual neural computations in humans that underlie the visual perception of bodies remain poorly understood. Most of the current evidence ^2–7^ comes from indirect measurements like fMRI, which offer limited resolution due to pooling activity across large neuronal populations and temporal averaging^8–10^. In contrast, single-unit data in non-human primates (NHPs, ^11^) provide rich insights into the neural mechanisms of body processing, but similar evidence in humans is entirely lacking^12^.

We wanted to tackle a number of specific questions about the neural representation of visually-presented bodies in humans. Are individual neurons in the extrastriate body area (EBA)^13^ selective for whole bodies, or body parts, and if the latter, are they selective for specific body parts (like hands)? Do body-selective responses arise very soon after stimulus presentation, as expected for a largely feed-forward process? How do we process more abstract depictions of bodies, such as outline drawings or stick figures? Are neural populations selectively responsive to photographs of bodies also selectively responsive to these stimuli? To that end, we used multi-unit spiking activity recorded from intracortical electrode arrays in or near the EBA in two neurosurgical patients undergoing refractory epilepsy monitoring.

The neuronal populations in this region exhibited a clear preference for human bodies over other visual stimuli, including faces, objects, and other body categories. Subsequently, we performed a series of experiments to comprehensively characterize the pattern of body-selective responses in the recorded arrays. Our results demonstrate a clear preference for human bodies, as well as selectivity for individual body parts. At the individual site level, the majority of body-selective sites coded for multiple body parts simultaneously, with distinct preferences for each body part. The body-selective representations within the recorded arrays extended well beyond photographic depictions of bodies, as predicted by previous fMRI work^13^. Overall, our findings provide the most direct evidence to date for body selectivity in the human extrastriate visual cortex, and start to reveal the neural representations and computations underlying this important component of social perception.

## Results

We performed microelectrode recordings (96-electrode Utah array) in the lateral occipitotemporal cortex (LOTC) of two patients with refractory epilepsy undergoing intracranial recordings for seizure detection (Figure 1a). CT/MRI co-registration revealed that both arrays were implanted in the lateral occipital cortex at MNI coordinates consistent with the Extrastriate Body Area (Figure 1b).

**Fig 1.**
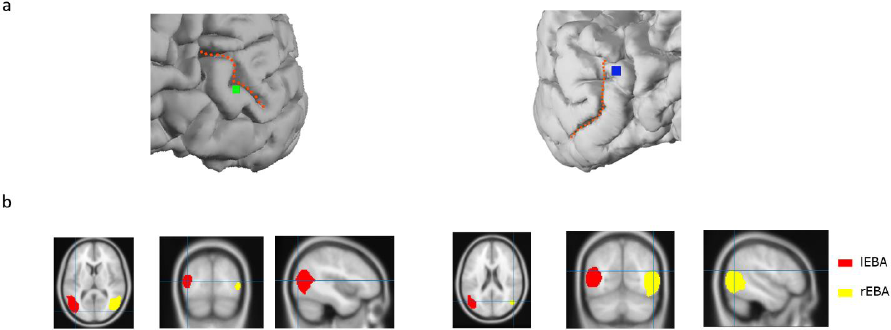
Multi-electrode array recording sites. **a**, Anatomical Utah array location on 3D rendering of the patient’s brain for P1 (Left, green square, MNI coordinates: X = -41, Y = -83, Z = 9) and P2 (Right, blue square, MNI coordinates: X = 51, Y = -66, Z = 19). Note the resection cavity above the array in P2. Dotted orange lines represent the Lateral Occipital Sulcus (LOS) **b,** Location of the Utah arrays relative to across-subjects EBA parcels identified in Julian et al^1^. Axial, coronal, and sagittal view for P1 (Left) and P2 (Right). Crosshair indicates the position of the Utah array for each patient.

### Category selectivity

To determine the visual responsiveness and underlying selectivity of the neural population, we first recorded the responses to 100 stimuli containing images of bodies, faces and other object categories (stimuli from Popivanov et al., 2012^14^, see Figure 2a for examples). We obtained data from 128 visually-responsive MUA sites across two subjects (two sessions in array P1 and one session in array P2, yielding 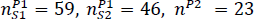 visually responsive sites, respectively).

**Fig 2.**
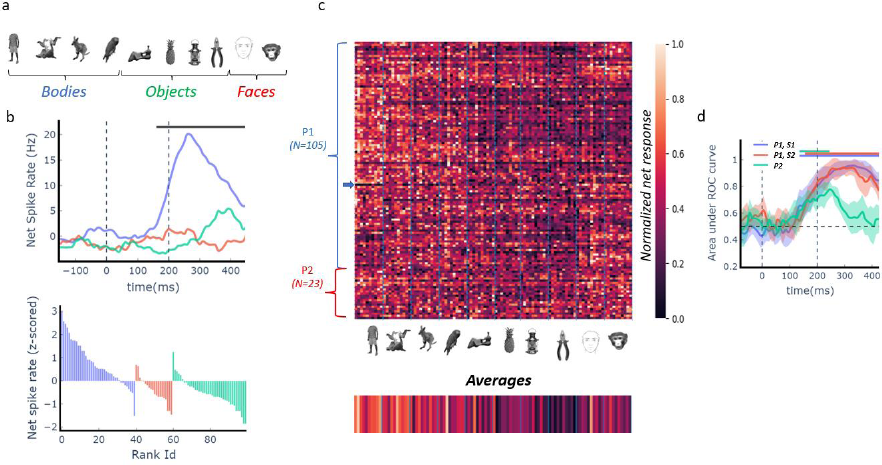
Category selectivity. **a**, Example stimuli from the category selectivity experiment grouped into Bodies (Blue), Objects (Green) and Faces (Red). **b,** Responses of a representative multi-unit example site. *Top panel* shows the time course of the mean net spike rate per category group. Horizontal bar represents significant response differences between categories (uncorrected Anova test per 100ms time point, sliding window of 10ms, P < 0.05). *Bottom panel*, average net z-scored responses to individual stimuli ranked within their respective category group (bodies, objects or faces). **c,** Color-coded normalized net responses of all visually responsive sites in array P1 (105 sites) and array P2 (23 sites) for the presented stimuli. Each row represents one recording site and each column represents one image. Columns were sorted by image category. **d,** Time course of decoder performance (area under the ROC curve; AU-ROC) for each recording session (blue: session 1 in P1; red: session 2 in P1; green: P2). Lines and shading show mean ± 1std classification AU-ROC for decoders trained to differentiate between stimuli containing a human body versus all other stimuli. Distribution reflects 10-fold grouped cross-validation. Horizontal bars indicate above-chance classification (P < 0.005; one-sided permutation test with at least five consecutive significant time points).

The response profile for an example recording site is shown in Figure 2b. The multi-unit activity (MUA) of this site responded significantly more to images of (human and animal) bodies than to images of (human and animal) faces or objects (one-way Kruskall Wallis ANOVA, P = 6e-24, Figure 2b, upper panel). For the majority of the body images, the responses were above the highest response to any face or object image (Figure 2b lower panel).

The population response selectivity was relatively heterogeneous (Figure 2c). On average, we observed a marked preference for images of bodies over other stimulus categories (see average responses in lower panel of Figure 2c). To quantify category selectivity for each individual site, we computed a category-selectivity index d’^15,16^ (see Materials and Methods). Significance was assessed by comparing the category d’ index to a null distribution generated by randomly permuting category labels across trials. More than half of the recording sites across the population were indeed body-selective (N=67/128 body-selective sites, permutation test, P < 0.05; 57.1% and 30.4% for array P1 and array P2 respectively, χ2 test, P < 4.17e-11, χ2=43.53). We also detected a smaller number of sites selective to faces (N = 27/128, permutation tests, P < 0.05, arrows in Figure 2b), but virtually no MUA site was object selective (N = 1/128, permutation test threshold, P < 0.05). Collectively, the observed body-selective responses and the anatomical location of the electrode recording sites (lateral occipital cortex) indicate that the recording arrays were likely located in or close to the extrastriate body area (EBA)^13^.

Next, we aimed to characterize the strength and time course of the population selectivity for human bodies in the recorded arrays. We employed linear decoders on the responses recorded in each session to assess the extent to which the pattern of neural activity elicited by a human body differed from that elicited by the other categories. Our decoding analyses were conducted in a time-resolved fashion on a trial-by-trial basis (see Materials and Methods for details). Specifically, we tested the neural pattern discriminability between human bodies and (1) the aggregate of the other categories (body vs non-body) and (2) the images within the body category (i.e., human bodies, monkey bodies, animals, and birds).

We found that a linear decoder trained on the population response could robustly discriminate between a human body and the other categories on a trial-by-trial basis in every recording session (permutation tests for each session P < 0.005; Figure 2d). Decoding performance was nearly optimal (AUC close to 1) in both recording sessions in P1, while the decoding in array P2 was less accurate presumably due to noise and/or the lower number of body-selective sites. The time courses of body selectivity were comparable across arrays in terms of the onset of selectivity (between 100 and 150 ms after stimulus onset) and the time of peak performance (250 ms after stimulus onset). This finding is consistent with the only other prior human invasive study examining body-selective responses around the EBA^12^ using subdural grid recordings (electrocorticography or ECoG). That study reported body-selective local field potential responses peaking at around 260 ms after stimulus onset in an ECoG contact at the approximate location of the EBA.

The recorded populations could reliably discriminate between different body subcategories with high accuracy, suggesting the presence of subordinate category representations (permutation tests P < 0.01; Supplementary Fig S1a). The confusion matrices associated with the subcategory-decoding (Supplementary Fig S1b) show that human bodies were the easiest to distinguish from the other subcategories. To confirm these findings, we trained individual decoders to discriminate between each different body subcategory in a one-vs-all fashion and found that decoders trained to discriminate human bodies versus all other body subcategories exhibited the best decoding performance in every recording session (Supplementary Fig S1c). To visualize the neural population representation, we performed a multidimensional scaling (MDS) analysis on the pairwise neuronal distances at the window of peak body selectivity (Methods). The MDS-derived two-dimensional neuronal object space (Supplementary Figure S2) further demonstrates that images of human bodies form a separate cluster distinct from the images of most other tested categories. Overall, our findings are in line with prior data that show body-selective areas of the human visual cortex showing a preference for human bodies^1^.

It is possible that all the results we report thus far were driven solely by the selectivity for low-level visual features. To address this concern, we compared the decoding results derived from our recorded neural data with those obtained using a state-of-the-art computational model of V1 activity (VOneBlock^17^, Methods). For both arrays (P1 and P2), human bodies were significantly easier to discriminate from the other categories using the recorded neural data compared to the V1 model (Supplementary Figure S3a, permutation test threshold, P < 0.001 for both sessions of array P1, P < 0.05 for array P2), suggesting that the neural selectivity for human bodies cannot be solely accounted for by low-level visual feature tuning. When further comparing the decoding performance for human bodies against the rest of the body subcategory images, the results remained statistically significant for array P1 (Supplementary Figure S3b, permutation test threshold, P < 0.05 and P < 0.001 for Session 1 and Session 2 of array P1 respectively), indicating a robust selectivity for human bodies over other body types in this array. However, for array P2, this within-category discrimination was higher for the neural data than for the V1 model but this difference did not reach statistical significance (Supplementary Figure S1d, p=0.18).

### Body selectivity is driven by features prevalent in human bodies

Our previous findings showed an overall preference for bodies over other tested stimulus categories. Next, we explored how the recorded populations responded to specific body parts. To this end, we presented a stimulus set that included images of whole bodies, isolated body parts (hands, feet, bodies without hands and feet, and heads), and images of objects (see examples in Figure 3a).

**Fig 3.**
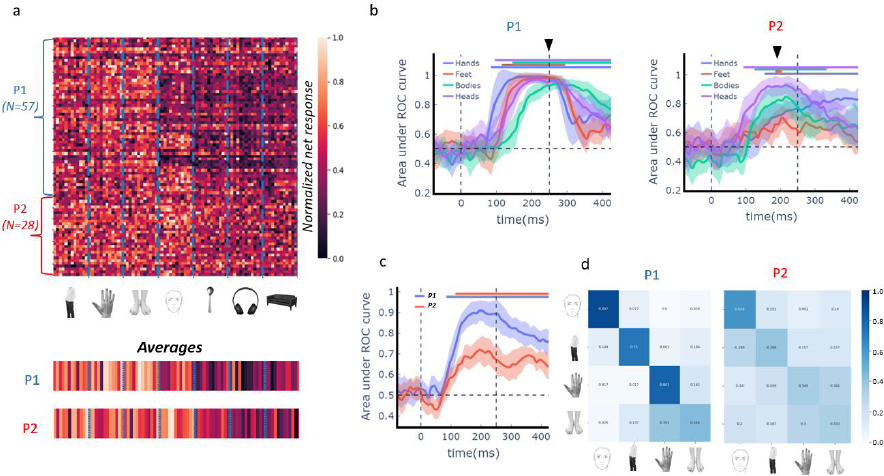
Body part selectivity. **a**, *Top panel* depicts the normalized responses of all visually responsive sites in P1 (blue, 57 sites) and P2 (red, 28 sites) to 84 images of whole bodies, isolated body parts (hands, feet, bodies without hands and feet, and heads), and images of objects (12 images per category, example stimuli shown in middle panel). Each row represents one site and each column represents one image. *Bottom panel* shows the average response to each image across all visually responsive sites in array P1 and array P2. **b,** Time course of performance of individual body part decoders (AU-ROC) for array P1 (left) and array P2 (right) trained to distinguish between individual body part and non-body images. Same conventions as in Figure 2d. **c,** Time course of performance of intra-body part decoders (AU-ROC) for both arrays trained to discriminate between different body parts. Same conventions as in Figure 2d. **d,** Confusion matrices for intra-body decoders at the peak decoding time window for array P1 (Left) and array P2 (Right). The x-axis represents ground-truth labels and the y-axis the predicted labels. The matrices are normalized across rows to highlight the proportion of true positives across each category.

We recorded 85 visually-responsive MUA sites in both subjects (one session in array P1 and one session in array P2, yielding *n^p1^* = 57, *n^p2^* = 28 visually-responsive sites, respectively). To quantify if the recorded neural population encoded body part-specific information, we trained linear decoders to (1) distinguish between individual body parts and non-body images (e.g., feet versus objects; “body part decoders”) and (2) discriminate between different body parts (“intra-body decoder”).

The body part decoders robustly discriminated between every individual body part and images of objects in both patients (permutation tests for each session P < 0.005; Fig 3b). The time course of decoding varied slightly for each body part and reached peak performance around 250ms after stimulus onset for array P1 and around 200ms for array P2, triangles in Figure 3c). Note that this time window also corresponds to the time at which human body selectivity emerged in the previous analysis (Figure 2d). We found near-optimal performance (area under ROC curve at perfect 1.0) for each individual body part in array P1, and a somewhat lower performance in array P2, in which the body-part information was dominated by a preference for heads and bodies (i.e. torsos with arms and legs) over hands and feet. Furthermore, the recorded neural population could reliably discriminate between different body parts (permutation tests for each session P < 0.005; Fig 3c) suggesting fine-grained body-part representations at the population level. This pattern was also more prominent in array P1 than in array P2 (see Figure 3d). The MDS-derived two-dimensional neuronal object space (Supplementary Figure S4) further confirmed our results since the images containing a body part (whole bodies, heads, hands, feet) were separable from images containing an object.

As before, when also tested these results against a CNN-based V1 model. Our results (Supplementary Figure S5) show that the fine-grained body-part representations emerged exclusively with our recorded data. For array P1, the body-part decoders trained using neural data significantly outperformed the V1 model across all body parts, while for array P2, all body parts except feet showed significantly better decoding, consistent with a preference for bodies and heads (Supplementary Figure S5a). Intra-body decoding also revealed superior performance for both patients compared to the V1 model (Supplementary Figure S5b). Taken together, these findings suggest that the body-selective section of the lateral occipital complex encodes specific body-part information that is linearly separable from non-body images.

Finally, we explored the representation of each body part by individual body-selective sites. We asked whether these sites were ‘selective’ or ‘general’ in their responses to different body parts. A selective site was defined as a site that had a significant d’ value (compared to objects) for only one specific body part (example selective single unit in Supplementary Fig S6), while a general site was defined as one that had a significant d’ value for more than one body part (example general single unit in Supplementary Fig S7). We found that the recorded populations consisted predominantly of general sites for both arrays (85.4% and 90.9% for array P1 and array P2 respectively, Supplementary Fig S8a). To further investigate how these general sites encoded different body parts, we examined the spatial distribution of the selectivity for each body part (hands, feet, bodies without hands and feet, and heads), with respect to objects, across both arrays (Supplementary Fig S8b). In array P1, we observed two distinct neural populations: one pool of highly selective sites for heads (left side of array P1) and another pool that responded to other body parts with different preferences. The results for array P2 indicate primarily body-selective sites with a preference for heads and bodies. Collectively, these results point towards a more diffuse code where body-selective sites encode multiple body parts simultaneously.

### Tolerance to shape-preserving image transformations

We tested the tolerance of the recorded neural representations to shape-preserving image transformations. In particular, we recorded responses to (1) silhouette versions of the category-bodies stimuli and to (2) bodies at different orientations.

Silhouettes lack internal details, which allows us to study the degree to which neural response selectivity depends on the external contours alone. We reasoned that if the body contour is a strong determinant in driving the responses of body selective neurons, intact images of bodies and body silhouettes would share a common population representation. To examine this, we tested whether decoders trained on intact body images (discriminating between images of human bodies and non-bodies) generalized to silhouette versions of the same images without retraining (Fig. 4a, Methods). We find that In both arrays, the trained body decoders could generalize from natural to silhouette versions of bodies, and vice versa. The decoding performance and observed time course for silhouettes was almost identical to the original decoders trained on intact images (Fig. 4b). Furthermore, in array P1, the generalization performance could not be attributed to tuning for low-level V1-like visual features (Figure 4c, permutation values, P < 0.001). Note that while array P2 exhibited higher generalization performance for neural data compared to the V1 model, this difference was not statistically significant (Figure 4c, P=0.11). Individual sites also generalized, representing intact and silhouette bodies similarly (Supplementary Figure S9). These results indicate a shared neural code for intact and silhouette body stimuli in the recorded population.

**Fig 4.**
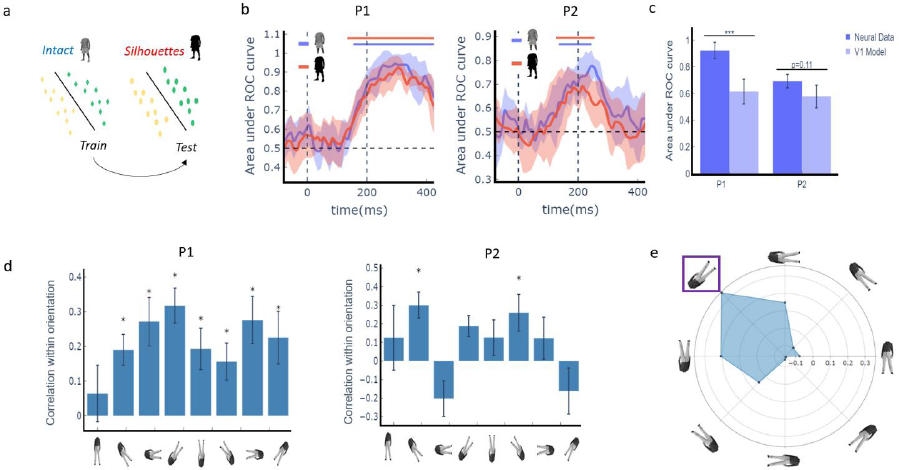
Tolerance to shape-preserving image transformations. **a**, Schematic illustration of decoders employed to investigate the shared population representation between intact images of bodies and their corresponding silhouettes. Body decoders were trained on intact images to discriminate between images of human bodies and non-bodies and tested on silhouette versions of the same images. **b,** Time course of body decoders’ performance (AU-ROC) for array P1 (left) and array P2 (right), for intact images (blue) and generalization decoders for silhouettes (red). Same conventions as in Figure 2d. **c,** Comparison of peak performance for generalization decoders trained with neural data compared to a model of V1-like activity. **d,** Average correlation of each population vector with itself (autocorrelation) across independent halves at each orientation for array P1 (Left) and array P2 (Right). **e,** Tolerance to body orientation in array P1. The polar plot illustrates the average correlation between each population vector at the orientation with the highest autocorrelation (purple square) and the population vectors corresponding to the same stimulus at different orientations. * P < 0.05, *** P < 0.001, one-sided permutation test

To explore tolerance to changes in body orientation, we computed the average correlation between each population vector at a reference orientation and the population vectors corresponding to the same stimulus at a different orientation (Methods). First, we examined the reliability of recorded populations for each orientation separately by computing the average correlation of each population vector at a particular orientation in half of the trials with the population vector in the other half of the trials. Significant information was present for all orientations except one in array P1 (permutation tests P < 0.05, Fig 4d left panel), while array P2 only showed significant information for two orientations (permutation tests P < 0.05, fig 4d right panel). This suggests that array P2 demonstrated limited tolerance to in-plane changes in orientation, as the available information was limited to a small number of orientations (45 degrees and 215 degrees) that were not spatially contiguous.

Next, we focused on array P1 to examine the similarity of the neural representations across orientations. The population of body-selective sites in array P1 showed tolerance to body orientation over a range of approximately 45 degrees (Fig 4e), yet the population representation changed markedly for larger orientation changes. Overall, our findings indicate that the responses of body-selective sites to images of bodies depend greatly on their orientation, highlighting the importance of orientation in shaping the neural representation of bodies.

### Representation of abstract bodies

Finally, we investigated how body-selective neural sites in human visual cortex encode highly abstract images of bodies. Stick figures are highly impoverished compared to naturalistic images of bodies, but humans have no problem correctly identifying and classifying these images as bodies. We compared the responses of stick figures with scrambled stick figures (to account for low-level visual features) together with outline drawings of objects (abstract fruits) and scrambled objects (example stimuli shown in Figure 5a). We recorded responses from 33 visually-responsive sites in both subjects (one session in array P1 and one session in array P2, yielding *n^p1^* = 13, *n^p2^* = 22 visually-responsive sites, respectively). The example site in Figure 5b responded significantly more (Kruskall-Wallis ANOVA, P < 5e-22) to images of stick bodies than to scrambled controls or fruits (Figure 5b, left panel) and, on average, the large majority of stick bodies evoked stronger responses than the scrambled bodies (Figure 5b, right panel).

**Fig 5.**
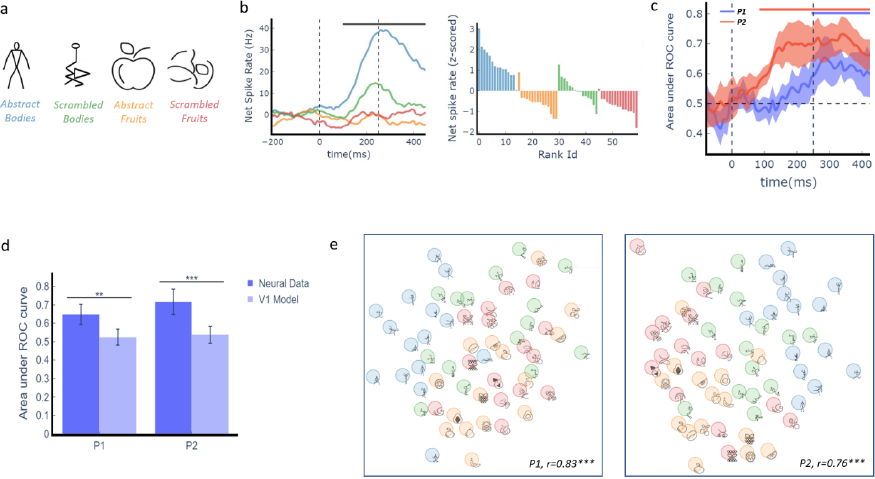
Responses to abstract bodies. **a**, Example stimuli from the different categories presented to assess abstract representation of bodies. **b,** Representative example site. *Left panel*, time course of mean net spike rate for abstract bodies (blue), scrambled bodies (green), abstract fruits (orange) and scrambled fruits (red). Horizontal bar represents significant response difference between categories (uncorrected Anova test per time point, P < 0.05). *Right panel*: averaged z-scored net responses to individual stimuli and ranked within their respective category. **c,** Time course of linear decoders performance (AU-ROC) for both arrays (P1 in blue and P2 in red). Lines and shading show mean ± 1std classification AU-ROC for each decoder trained to discriminate the category of each stimulus. Same conventions as in Figure 2d. **d,** Comparison of peak performance for decoders trained with neural data (c) compared to a model of V1-like activity. **e**, Visualization of similarity relations in the population evoked response for the presented stimuli obtained using multidimensional scaling (MDS) for P1 (left) and P2 (right). ** P < 0.01, *** P < 0.001, one-sided uncorrected permutation test.

At the population level, linear decoders trained to distinguish among the four stimulus classes performed significantly above chance for both patients (corrected permutation tests P < 0.005, Figure 5c). This performance cannot be explained by tuning for low-level visual features (Figure 5c, permutation test, P < 0.01 and P < 0.001 for arrays P1 and P2, respectively). The confusion matrices (Supplementary Figure S10) show that stick bodies were more discriminable than other stimulus categories, which is also visualized in the associated MDS (Fig. 5d) where the intact stick figure cluster (blue) is clearly distinct from their scrambled counterparts (green) and intact and scrambled fruit-like images.

These findings suggest that the recorded populations respond preferentially to bodies even when the images are reduced to stick figures. However, this result does not address the question of whether the neural representation of stick bodies overlaps with the neural representation of full-cue presentations of human bodies. To explore this further, we presented a reduced version of the original EBA set localizer of Downing et al. 2001, which contains different bodies and body parts together with stick figures of bodies (only one session in array P2, yielding 32 visually-responsive sites; see Methods and Materials). We first confirmed our previous findings by showing that every individual body category was linearly separable from the other object categories (corrected permutation tests P < 0.005, Supplementary Fig S11a), including images of bodies when presented as stick figures (corrected permutation tests P < 0.005, Supplementary Fig S11b). Next, we performed a decoding generalization analysis analogous to the silhouette experiment. Specifically, we tested whether linear decoders trained to discriminate between images of a bodies and objects could generalize when tested with stick figures (intact versus scrambled). We observed partial but significant generalization as the ability of the classifier to generalize in this way reached significance but was considerably reduced compared to the original classifier (Supplementary Fig S11c). Moreover, the generalization performance was significantly higher with the recorded neural data compared to V1-like activity (Supplementary Fig S12b, permutation test, P < 0.01). These results suggest partially overlapping population representations of human bodies and stick figure bodies. Consistent with this idea, the two dimensional MDS neural representation showed separate clusters for bodies and objects, with stick figure bodies placed in between (Supplementary Figure S11e).

## Discussion

Here we presented a series of experiments to characterize body-selective responses in the human lateral occipital cortex at the level of multi-unit activity. Our results show that the neural responses in human body selective cortex exhibit a clear preference for human bodies over other visual stimuli tested (including faces, objects, and other body categories), are driven by individual body parts, show orientation dependence, and generalize from naturalistic (photographic) images of human bodies to highly symbolic/abstract images of bodies. The spatial and temporal resolution of the reported body selective responses (multi-unit spiking activity) is unprecedented for studies in humans and sheds new light on the neural mechanisms supporting the visual perception of bodies.

The observed pattern of body-selectivity and the implantation sites of our electrode arrays in lateral occipital cortex strongly suggest that the Utah arrays recorded from the extrastriate body area (EBA)^13^, a body selective region broadly implicated in human body perception^1,18^. While body-selectivity has been studied extensively using functional MRI, this is the first direct evidence for strong body selectivity in humans at the level of multi-unit spiking activity (Pourtois et al^12^ for a single electrode ECoG array). Few prior fMRI studies^3,19^ also suggested neural sensitivity for body parts in the human EBA, but since fMRI measures the pooled activity of multiple thousands of cortical neurons, it remained unclear whether selectivity for individual body parts would be observed at the neuronal level in humans. Our study provides the strongest empirical support for the existence of a body selective neural population tuned to discriminate between individual human body parts. We also observed non-trivial differences in the preferences for different body parts across the recorded arrays. In particular, array P1 (located more ventrally) had more decodable information about each body part (with higher responses overall for hands and feet, which agrees with previous fMRI studies^20,21^), while the more dorsally located array P2 showed a clear preference for heads and bodies (without head, hands and feet). Orlov et al.,^22^ previously used reverse correlation analyses to suggest a mosaic of regions (encompassing the EBA) that respond in subtle but systematic ways to different body parts. Qualitative comparisons suggest that array P1 likely recorded from the area previously reported as the “upper-limbs” region (which mostly overlaps with the EBA), while array P2 may have been located in the “upper-face” area. We also observed that responses in array P1 emerged faster and were stronger compared to array P2. Despite these differences, all the main findings of this study (body and body part selectivity, contour selectivity, orientation tolerance, representation of stick figures) were replicated in both patients.

We further report that the body-selective sites encode multiple body parts simultaneously, with distinct preferences for each body part. This suggests that body representations are not based on semantic labels such as “finger”, “hand” or “foot”, but on the similarity between visual features. We interpret this finding as a neuronal correlate of “cognitively unelaborate” body representations consistent with the proposal in Downing and Peelen^23^. The neural responses in this region are driven by the visual shape of body parts, not by their high-level semantic labels. Conversely, the features that drive the recorded populations are more complex than simple low-level features. This becomes evident when we compare our decoding analyses with a state-of-the-art computational model emulating V1 activity. Most likely, the lower decoding performance in array P2 was due to lower signal to noise in array P2 recording electrodes as well as fewer body-selective channels (see Supplementary Fig S13). Both factors are known to lower the ability to decode information from the neuronal population. Critically, all the decoding trends in array P2 were in line with the findings in array P1.

Downing et al.^13^ previously showed that the body-selective representations within the EBA may generalize to other abstractions such as silhouettes, line drawings, and stick figures. Our results indicate that the degree of generalization at the neuronal level is not the same for different kinds of abstractions. On the one hand, we found that the response selectivity generalizes strongly from intact bodies to silhouettes, which provides support for the notion that shape features drive the selectivity of neurons in higher-level visual cortex^24,25^. In contrast, the results for stick figures indicate that while they are preferentially represented compared to non-body stimuli, their neural representation only partially aligns with the neural representation of photorealistic bodies. Notably, this partial generalization was absent when we considered their scrambled counterparts, suggesting that the preferential responses observed for the stick figures cannot be explained by selectivity for low-level features.

Body selectivity has also been observed in the macaque extrastriate cortex ^14,26^. The macaque mid-STS body (MSB) patch was suggested as the homologue of the human extrastriate body area (EBA), even though neural properties had never been formally compared. Our findings reveal a striking similarity between the properties of human body-selective neurons and those found in the MSB^27^. Notably, we observed similar selectivity for intact and silhouette bodies consistent with Popivanov et al^28^, and the recorded neurons exhibited a limited tolerance to image rotation, with strong responses to specific body parts, as in Popivanov et al^14^. Additionally, population-level analyses demonstrated accurate decoding of human versus monkey bodies, while faces or heads were distinguishable from bodies and body parts in line with Kumar and Vogels^29^. These findings provide crucial evidence for the macaque monkey as a valid model for studying body processing in the visual system. Future studies will have to include intracortical recordings in other nodes of the human body processing network to chart the homology between the human and macaque body processing systems.

Further insights into the neural underpinnings of body perception will require an end-to-end mechanistic understanding of the specific visual features that drive the selectivity of body selective neurons^23^. One possible approach is the use of computational models. Deep neural networks have been used to successfully predict the responses of single neurons in macaques^30–33^ and fMRI bold activations in humans^34,35^, including category-selective regions like the EBA^36,37^. High-resolution neural data from the human EBA, as in this study, can be used not only as a strong experimental constraint for computational models, but also as a tool to uncover the specific visual features that drive responses of human cortical neurons (for example using artificially synthesized stimuli, Ratan Murty et al^36^). There are several ways to extend our work using such approaches. First, our data offer the possibility of developing a model per recording site, which can be compared against previous fMRI-based pooled models of the entire EBA. Second, our data provide ground-truth responses to images that fall well outside the manifold of naturalistic images used in previous studies (such as silhouettes and abstract bodies). Whether or not predictions from naturalistic image generalize to stimuli outside the training distribution remains an open question^38,39^. Third, electrophysiological data in our study are dynamic and the temporal resolution of MUA recordings provides an additional constraint for (generally feedforward) models to explain^40^. Together, the synergy between computational approaches and high-resolution spiking neuronal activity recordings from human EBA promises to uncover the neural mechanisms of body perception in humans at a finer grained and quantitative level.

## Methods

### Study approval

We obtained ethical approval (study number s53126) for conducting microelectrode recordings using the Utah array in patients with epilepsy. The study protocol (s53126) was approved by the local ethical committee (Ethische Commissie Onderzoek UZ/KU Leuven). The study was carried out in accordance with the principles of the Declaration of Helsinki, the principles of good clinical practice, and all applicable regulatory requirements set by the Federal Agency for Medicines and Health Products (FAGG).

To ensure safety and compliance, strict adherence to imposed safety measures was maintained. This included the use of case report forms and detailed reports on any (serious) adverse events. All data collected during the study were encrypted and stored at the University Hospitals Leuven. The study was discussed with the patients during the pre-operative consultation, which took place more than six weeks before the surgery. The patients were informed about the additional risks associated with micro-electrode array implantation, such as infection and local hemorrhage. Written informed consent was obtained from the patients on the evening prior to their surgery.

### Clinical information

We obtained invasive intracranial recordings from the lateral occipital cortex in 2 refractory epilepsy patients. Patient 1 was a 58-year-old woman who had never undergone brain surgery. Patient 2 was a 29-year old male who had undergone resective epilepsy surgery from the right temporoparietal junction at the age of 19 (MNI coordinates anterior border (64, -42, 32), superior border (51,-53,56), posterior border (55,-67,42) and inferior border (60,-60,27)). The pathology report after this resection mentioned chronic meningitis. After a year of seizure freedom, his seizures recurred, and invasive intracranial recordings were performed as diagnostic epilepsy surgery workup. Patient 1 was on Levetiracetam 2000 mg, Diphantoine 100 mg, Rivotril 1 mg and Lacosamide 300 mg, all taken twice daily. Patient 2 was on Levetiracetam 2000 mg twice daily and Lacosamide 300 mg twice daily. During hospitalization, medication was adjusted in both patients according to the number of detected seizures. Implantation sites were left lateral occipital cortex (MNI coordinates -41, -83,9) in patient 1 and right lateral occipital cortex in patient 2 (MNI coordinates 51,-66,19). Although the presumed epileptogenic zone (PEZ) encompassed the lateral occipital cortex based on multimodal preoperative assessment, invasive recordings showed no clear focal epileptic onset in patient 1. In patient 2, seizures originated from the anterior border of the previous resection, and a new resection was performed at the time of electrode removal. In both patients, the array was outside the actual epileptogenic zone. The multi-unit spiking activity on the arrays was analyzed to investigate to what extent neural activity could predict seizure onset (“Microscale Dynamics of Epileptic Networks: Insights from Multiunit Activity analysis in neurosurgical patients with refractory epilepsy”, Bougou et al., EANS 2023, Barcelona).

Target locations of intracranial arrays were determined by the epileptologist and were based on electroclinical findings and non-invasive multimodal imaging. The multielectrode arrays were implanted via the craniotomy performed for the implantation of subdural grids and were thus inserted in close proximity to this grid. No additional incisions were made specifically for the purpose of this study. The arrays were removed during the same surgery for subdural grid removal 14 days after implantation.

### Invasive recordings

We recorded from a 96-electrode array (Utah ArrayTM – Blackrock Neurotech) from day 1 until day 14 after array implantation. We inserted the array using a pneumatic inserter wand provided by Blackrock Neurotech. We closed the dura above the array and placed the bone flap on top to secure the array. We placed reference wires subdurally and ground wires epidurally. We digitally amplified the signal using a Cereplex M head stage, connected to a 128-channel neural signal processor (Blackrock Neurotech), and sampled the signal using Central Software at a sampling rate of 30 kHz. We applied a 750 Hz high-pass filter for spiking activity. We set the multiunit detection threshold at 3 standard deviations of the noise band. We considered units recorded on the same electrode but on different days as separate units. The variable signal quality in the immediate postoperative period resulted in a difference in the number of responsive and selective sites during each recording session. All spike sorting was performed offline (Offline Sorter 4, Plexon, TX).

### Stimulus presentation

We conducted experiments in a dimmed hospital room. We utilized custom-built software on a 60 Hz DELL-P2418HZM LED monitor to present stimuli. Patients sat 60 cm away from the screen (1 pixel = 0.026). We instructed them to focus on a small red square (0.2 × 0.2°) positioned at the center of the display. We continuously monitored the position of their left or right pupil using a dedicated eye tracker (Eyelink 1000 Plus, 1000 Hz) in head free-to-move mode. If they deviated from the electronically defined 3° by 3° fixation window, the trial was aborted. To synchronize the data, we attached a photodiode to the upper left corner of the screen, which detected a bright flash coinciding with the first frame of the stimulus. This flash was invisible to the patients. To accurately record baseline spike rate activity, we introduced an intertrial interval between the offset of the stimulus and the onset of fixation for the next trial. Additionally, to maintain fixation, we instructed the patients to press a button with their right hand whenever a distractor (red or green cross) randomly appeared at the fixation point in approximately 2% of the trials.

### Stimuli

#### Category-bodies

The stimulus set consisted of 100 achromatic stimuli grouped over 10 categories (Popivanov et al, 2012^14^), each stimulus was presented ∼7-10 times. Categories included mammals, birds, fruits, bodies (human–monkey), faces (human–monkey), objects (matched to the aspect ratio of images from the human class “objectsH” and monkey class ”objectsM”, respectively) and sculptures. The mean luminance values were equated across classes. The mean vertical and horizontal extent of the images was 6.0° and 4.6°. Stimuli were presented on a uniform gray background. After a fixed fixation period of 300 ms, stimuli were presented for 200ms with an intertrial interval of 150ms.

#### Silhouette tolerance

We presented silhouette versions of every stimulus from the category-bodies set randomly interleaved with stimuli from the categorical set in one session for each patient. Each stimulus was presented ∼7-10 times. The size of the silhouette version of every stimuli corresponded to their intact counterpart. Stimuli were presented on a uniform gray background.

#### Body part responses

We presented a stimulus set that included images of bodies and isolated body parts as well as images of objects. The images from every category except for the feet categories are from the set employed in Matic, et al., 2020^41^. The included conditions from this set comprised three categories of different body parts: bodies (only torso; without the head, hands, and feet), hands, and heads and three object categories validated with behavioural judgements (i.e., tools, manipulable objects, and nonmanipulable objects). The stimuli from the different object categories were assembled such that category information could not be extracted based on their physical shape. All images were controlled for low-level visual properties (e.g., spatial frequencies and luminance) by means of the SHINE toolbox^42^. Each condition consisted of 12 achromatic images ( ∼12 repetitions per stimulus) on a white background (Fig. 3A). The mean vertical and horizontal extent of the images was 8.3° and 8.1°.

#### Tolerance in-plane body rotations

Four bodies from the category selectivity task were shown within the receptive field at 8 different rotations (45° angles between each rotation, starting from upright position). Body size was 4°. In patient 1, stimulus location was at (6°,3.5°) in the lower contralateral quadrant, in patient 2 at (0°,0°). Each stimulus was repeated ∼12 times for every rotation. Patients performed a distractor task as described above.

#### Abstract body representations

To investigate how body-selective neurons responded to abstract representations of human bodies, we utilized stick figures as abstract representations, while also employing scrambled versions of these stick figures as a control to account for low-level visual features. Stimuli consisted of 60 black line drawings on a white background grouped in 4 categories (bodies, scrambled bodies, fruits and scrambled fruits). Each body and fruit image were matched to 1 scrambled stimulus which contained the same line fragments. Scrambled counterparts were manually made by reorganizing the individual lines composing the original body or fruit. The mean vertical and horizontal extent of the images was 6.6° and 6.0° . Patients performed a distractor task as described above (∼ 2% of trials).

#### Reduced EBA localizer

The stimulus set consisted of 125 achromatic stimuli grouped over 6 categories, each image was presented ∼7-10 times. Categories included whole bodies (photographic and drawings), body parts (photographic and drawings), objects (photographic and drawings), places, stick figure bodies and scrambled stick figures. The images from every category except the place category come from the original EBA localizer of Downing et al., 2001^13^. Stimuli were presented on a uniform white background. The mean vertical and horizontal extent of the images was 6.8° and 6.3° of visual angle. Patients performed a distractor task as described above (∼ 2% of trials). Fixation duration was 300ms, stimulus duration 250ms and intertrial interval 150ms.

### Imaging

We acquired a T1 weighted image using a 3-T MR scanner (Achieva dstream, Philips Medical Systems, Best, the Netherlands) at the University Hospitals Leuven for presurgical planning and to reconstruct the postoperative electrode location. The imaging parameters were as follows: time of echo 3.1 ms, inversion time 900 ms, slice thickness 1.0 mm, 400 × 400 matrix, and 283 coronal slices. Following electrode placement, a CT scan (Siemens) was performed with a slice thickness of 1 mm, a voltage of 120 kV, and a dose length product of 819 mGy.cm. The CT scan served to verify the electrode locations and to rule out any hemorrhage.

For imaging preprocessing, we utilized SPM12 software (Wellcome Department of Cognitive Neurology, London, UK) running on MATLAB (Mathworks, Natick, MA). The preprocessing steps included (1) Setting the anterior commissure as the origin for both the MRI and CT scan (2) Realigning the images (3) Coregistering the MRI and CT images and (4) Warping the MRI and coregistered CT to a common high-resolution brain atlas (MNI1525^43^). To generate cortical 3D renderings of the subjects’ pial surfaces and the high-resolution MNI atlas, we employed Freesurfer. The precise electrode location on the cortical surface and the MNI coordinates were determined using iElectrodes Software^44^.

### Single-site data analysis

We computed multi-unit firing rates for each correct trial in two analysis windows: a baseline window ranging from 150 to 50 ms before stimulus onset and a response window ranging from 200 to 300 ms after stimulus onset. We restricted all analyses to visually responsive sites. To assess the responsiveness of each recorded site, we performed independent t-tests between the spike rates in the baseline and response windows for each stimulus category separately (e.g. baseline vs all human bodies). For every experiment separately, we considered a site as visually responsive if any of these comparisons was significant (P < 0.01). The reasoning behind this criterion is that sites with a strong preference for a single category could have been considered as not responsive when testing baseline vs response across all stimuli in the test. All subsequent analyses were based on the baseline subtracted, average net firing rates. In cases where responses across sites were shown on the same figure, we scaled the average net firing rates for each site in the range (*x_min_* = 0, *x_max_* = 1) as follows:

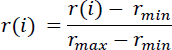

In which, *r*(*i*) corresponds to the average net response for stimulus *i* for that site *r*_*min*_ and *r*_*max*_ correspond to the “best” and “worst” stimuli for that site, respectively.

#### Calculation of d’

To quantify category selectivity for individual sites, we computed d prime index or d’^15,16^, which takes into account the mean responses to the corresponding categories as well as the variability of the responses to the individual images of a category. The d’ indices were computed for different pairs of categories using the net evoked firing rates. Here, we illustrate the computation of d’ in the example case of comparing trials in which an image of a body or non-body was presented:

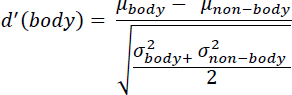

In which 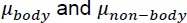 are the mean firing rates and 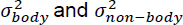 are the variance in firing rates on trials in which an image of a body or non-body was presented, respectively. For this analysis, we removed the responses to images from the birds and mammals categories as they included both a body and a face. The significance of each site’s d’ was assessed by comparing the values obtained to a null distribution of values generated by randomly permuting category labels (body or non-body) across trials (1000 iterations).

#### Noise-ceiling estimates

The noise-ceiling serves as a measure of the reliability of the recorded responses, taking into account the inherent noise in the data. To obtain these estimates, we randomly split the trials into two groups, repeating this randomization 100 times. For each experiment, we estimated the noise-ceiling of every site by first computing average normalized net spike rate per image separately for each half of the data, and then measuring the (Spearman-Brown corrected) correlation between the average responses from the two halves. This entire procedure was repeated 100 times for each binary split.

### Population Analysis

#### Multidimensional scaling

To visualize the similarity relationships between objects in the neuronal population, we employed multidimensional scaling (MDS). The neuronal distances were computed as the average Euclidean distance of the firing rates across neurons for each image pair (alternative distance metrics such as correlation or cosine distance yielded comparable results). We utilized the MDS implementation from the scikit-learn library (implemented in Python). In line with our single-site analysis, we utilized the baseline-subtracted average net spike rates across repetitions of the same image. To ensure equal contribution of each site to the data variance, spike counts were z-standardized.

The MDS procedure was applied to the response matrix of each experiment, with dimensions corresponding to the number of visually responsive sites by the number of individual images. To evaluate the alignment between the two-dimensional embedding and the actual neuronal distances, we calculated the correlation between them. The resulting correlation coefficient is displayed in each multidimensional scaling plot. Notably, the observed high correlation values indicate that the multidimensional scaling plots provide a reasonable approximation of the underlying similarity relationships.

#### Decoding category information

We used linear decoders to quantify the amount of information about different stimulus categories in the recorded population of neurons on a trial-by-trial basis. We performed this analysis separately for each recording session using every recorded site to prevent bias towards any category. To quantify neural information in a time-resolved manner, firing rates were calculated within 150 ms bins with a sliding window of 10 ms. For each time bin, we trained a logistic regression classifier with L1 regularization and C=0.1, implemented using the scikit-learn library, to discriminate between categories based on the population firing rate at that time bin. The specific categories chosen depended on the experiment and hypothesis being tested. To address class imbalance, we adjusted the logistic regression loss function weights inversely proportional to class frequencies in the input data. For the multiclass problems (e.g. Figure 3d), we trained the decoders using a standard “one-vs-rest” approach. Specifically, a linear model was trained to discriminate between each individual category and the remaining categories in a binary fashion. Then, the population response in a specific time bin (the sample) was fed to every individual model and the sample category was attributed to the corresponding category of the model with the higher probability output.

To evaluate the decoding performance, we used the area under the ROC curve metric (AUC), which is insensitive to class imbalance as it is based on the true positive and false positive rates rather than the actual number of instances in each class. Classification AUC was estimated using a 10-fold exemplar-grouped cross-validation approach. In each validation step, we split the trials so that each repetition of a particular image could be in either the training or the test set, with each image exemplar being tested at least once. Before classification, net spike counts were z-standardized to ensure that classification was not influenced by the absolute magnitude of the responses. Training and test sets were not scaled independently but the same scaling was applied to both. To statistically compare decoding AUC to chance performance we computed for each time window the null AUC distribution by shuffling trial indices. For each time window, shuffling was repeated 250 times. Subsequently, actual decoding performance was compared to the null distribution using a one-sided permutation test and by considering only clusters with 5 consecutive significant bins.

#### Correlation of stimulus representations under shape-preserving image transformations

We wanted to assess the tolerance of the recorded populations to different identity-preserving image transformations in a time-resolved manner. We reasoned that if the neuronal population is tolerant to a specific transformation (e.g., 2D plane rotation) then the representation for each stimulus at the reference condition and the transformed condition should be correlated. To investigate this, we computed the average correlation between each population vector at a reference orientation and the population vectors corresponding to the same stimulus at a different orientation.

In order to provide reliability of the data given the noise and to provide confidence intervals, we binned the data according to each respective transformation and then randomly partitioned the data into two halves. In particular, trials were grouped into S = number of stimuli (varies over experiments), T = 2 pair of transformations (e.g. rotations of 45degs and 90degs), and H = 2 halves, yielding S × T × H = M total conditions. For each of these conditions, at the evoked response period, we computed the average population vector *r_S,T,H_*. Finally, the average correlation of the population response across transformations was computed as follows:

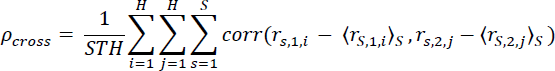

Where 〈∗〉_*x*_ corresponds to the average response vector across the set of stimuli S. Namely, for each set of S population vectors corresponding to a particular half of the data H and transformation T, we subtracted the mean across transformations to center the population vectors around zero. Therefore, *ρ_cross_* measures to what extent stimulus population representations are similarly organized around their mean across the two transformations.

We computed the average correlation of each population vector with itself across the two halves of the trials with the aim of obtaining an upper bound to *ρ*_*cross*_ given the degree of noise in the data:

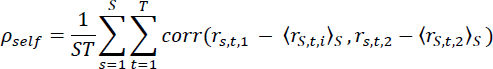

This analysis included only sites that responded significantly to at least one of the conditions (or rotations).

To obtain confidence intervals for *ρ*_*cross*_ and *ρ*_*self*_, we randomly split the trials for each condition (stimulus–transformation) into two groups, repeating this randomization 1000 times. The average firing rates across these sampled trials provided the mean population vector for that condition on that iteration.

To determine the significance of the computed *ρ*_*cross*_ and *ρ*_*self*_ values, we conducted permutation tests. In these tests, we shuffled the labels of the trials associated with the analyzed transformations (reference and target) to create a null distribution. For each iteration of the permutation test, we followed the same resampling procedure described earlier to obtain an average metric value for that iteration. Specifically, we performed 1000 permutation iterations, with 1000 resampling iterations per permutation iteration.

### Decoding generalization of category information under certain image transformations

We conducted a generalization decoding analysis to investigate the ability of the recorded neuronal populations to encode category information despite certain image transformations. We reasoned that if the population is tolerant to certain image transformations, responses to intact and transformed images would share a common population representation. In such case, a decoder trained on intact (original) images should exhibit significant classification performance when tested with transformed stimuli.

To evaluate the decoding performance for silhouette images, we performed a cross-validation training procedure where we trained the decoders using a subset of the intact images and we tested on held-out images for both intact images and silhouette versions; the held-out images belonged to the same object exemplars for both intact and silhouette images. In the case of decoding generalization in the abstract body test (Supplementary Fig 8), the images from the intact bodies and the abstract body images came from entirely different distributions. To assess significance for the decoding performance of the transformed sets, we performed permutation tests where the null distribution was created by shuffling the labels of both the training set and the generalization test set.

### V1 model control

To investigate whether the observed decoding results could arise from neural activity tuned for low-level visual features, we employed a V1 model (VOneBlock^17^) consisting of Gabor filters. In brief, the VOneBlock model is based on the classical linear-nonlinear-Poisson (LNP) model, consisting of a Gabor filter bank (GFB), simple and complex cell nonlinearities, and Poisson-like stochasticity.

To simulate a V1-like dataset for each experiment, we passed the stimuli through the V1 model. To account for the variability in the recorded neural responses, we simulated the repetitions of single trials by passing the same image multiple times, equal to the number of repetitions per stimulus in the original experiment. As the simulated data was subject to Poisson-like noise, we obtained different activation patterns for the same image, resembling the statistical characteristics of real neural data. The model convolved the input images with 512 Gabor filters, each having different orientations, sizes, shapes, and spatial frequencies. This process generated a 56x56 spatial map for each image, resulting in a stack of 512 output images, one for each type of Gabor filter.

To ensure a fair comparison between the neural data and the model results, we performed a subsampling procedure. We repeated this process 1000 times, randomly selecting a number of V1 units equal to the visually responsive sites specific to the array and experiment. For each iteration, we assessed decoding performance using group cross-validated linear decoding, following the same procedure as in the neural data analysis. Finally, we assessed the significance of the comparison between neural data and the V1 model results by calculating p-values as follows:

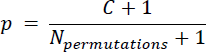

Where C is the number of permutations whose performance was higher than the one obtained with neural data.

## Funding

This work was supported by Fonds Wetenschappelijk Onderzoek (FWO) grant G.0B6422N, KU Leuven grant C14/22/134, and HBP SGA3 945539. T.T. is supported by FWO (senior clinical researcher; FWO 1830717N). M.V. holds a personal FWO grant for fundamental research (1169321N).

## Author contributions

We thank both patients for participation in this study. We thank Stijn Verstraeten for technical assistance for the recording setup and Anaïs Van Hoylandt for all help provided in data recording and data monitoring.

T.T. and M.V. planned and performed array placement surgery. M.V. performed the recordings and was responsible for all clinical trial related activities. J.G.R. performed the data analysis and wrote the manuscript. T.T. and P.J. supervised and guided the study. W.V.P. selected the patients for invasive brain recordings. All authors reviewed and edited the manuscript.

## Declaration of Interests

The authors declare no competing interests.

## Supplementary figures

**Supplementary Fig S1.**
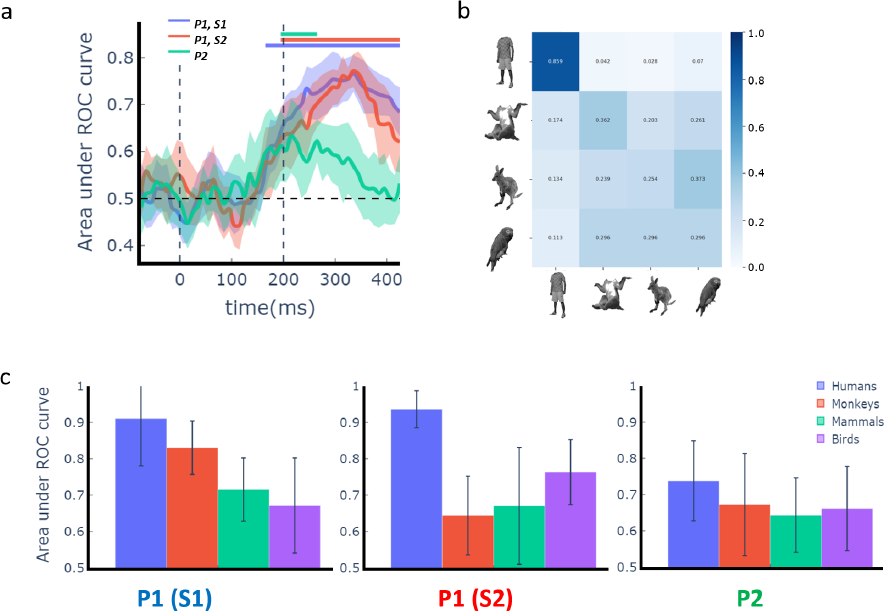
Neural information within the body category. **a**, Time course of decoder performance (area under the ROC curve; AU-ROC) for each recording session (labelled in top right). Decoders were trained to discriminate stimulus categories using images that exclusively contain a body (human, monkey, animals, birds). Lines and shading show mean ± 1std classification AU-ROC. Distribution reflects 10-fold grouped cross-validation. Horizontal bars indicate above-chance classification (P < 0.005; one-sided permutation test with at least five consecutive significant time points). **b,** Confusion matrix for within-body decoding during the peak decoding window for one session from array P1. The x-axis represents predicted labels, while the y-axis signifies ground-truth labels. Matrices are row-normalized to emphasize the proportion of true positives within each category. **c,** Comparison of performance for individual decoders trained to discriminate between each different body category in a one-vs-all fashion (e.g. human bodies versus rest of body images). Decoders were trained using activity within the evoked window of each array (200 to 300 ms after stimulus onset). **S1: session 1; S2: session 2.**

**Supplementary Fig S2.**
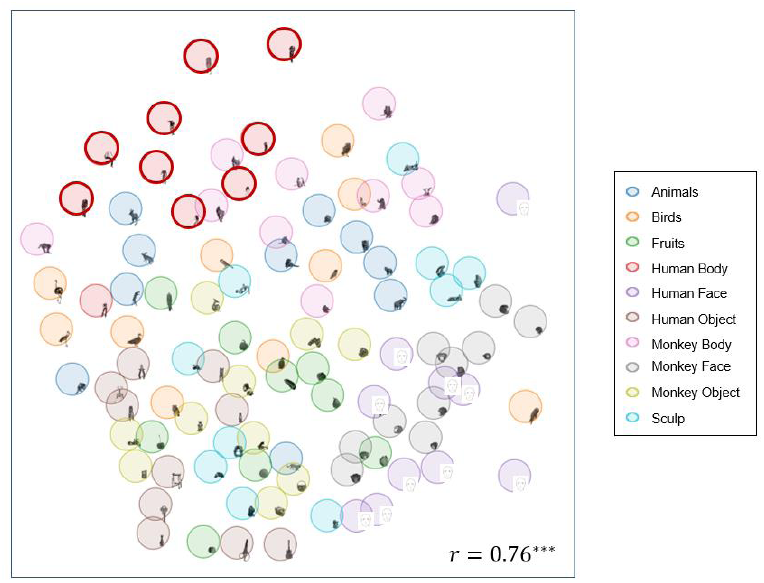
Similarity relations in the population response for categorical-body stimuli. Visualization of similarity relations in the population response to the categorical-body stimuli as obtained using multidimensional scaling (MDS). Evoked responses (200 to 300 ms after stimulus onset) from visually responsive sites (105 sites for array P1 and 23 sites for array P2) were utilized for the MDS plot. Proximity of images in the plot indicates similar neural response patterns across the population. The high Pearson correlation (r=0.76) between distances in neural space and distances in MDS space indicates that the presented two-dimensional MDS plot is a reasonable approximation to the observed similarity relations. *** P < 0.001, Pearson correlation between distances in neural space and distances in MDS space.

**Supplementary Fig S3.**
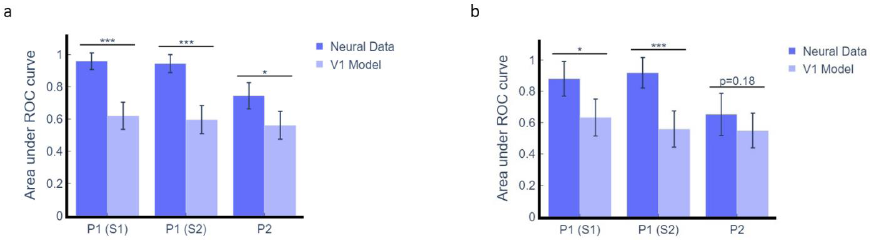
Comparing category information of neural data with V1 control model. **a**, Performance comparison between decoder trained with neural data to discriminate human bodies from all other categories (Fig 2d) and its site-matched model employing V1-like activity. **b** Performance comparison between decoder trained with neural data to discriminate human bodies from the rest of body categories – monkey, mammals and birds - (blue bars in Supplementary Fig S1c) and its site-matched model employing V1-like activity. Distribution for each V1 model represents 1000 iterations of randomly selecting a number of V1 units equal to the visually responsive sites of each recording session. * P < 0.05, ** P < 0.01, *** P <0.001 one-sided uncorrected permutation test.

**Supplementary Fig S4.**
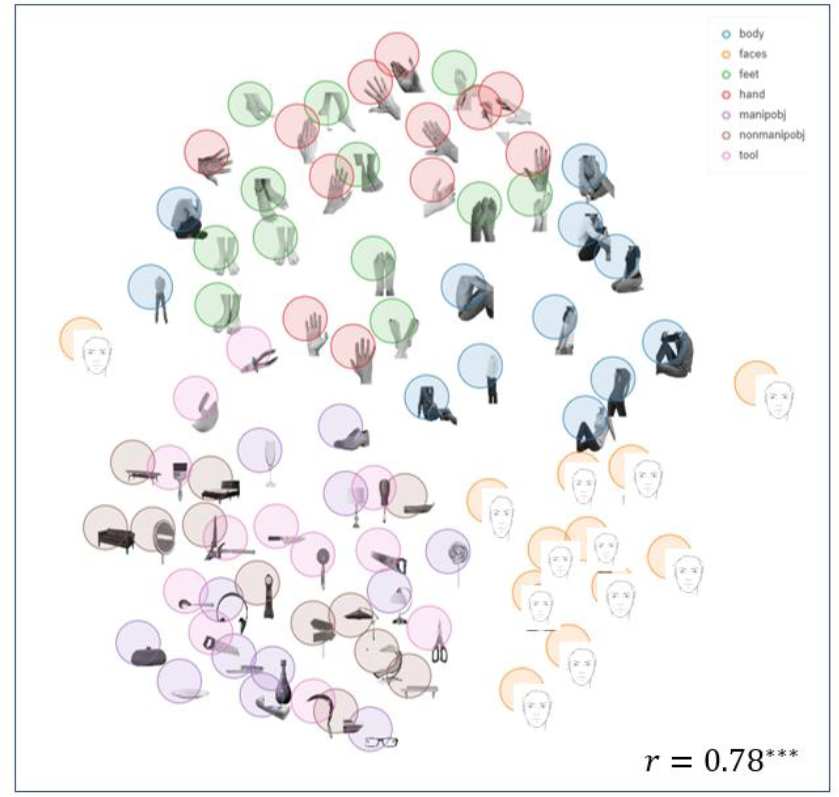
Similarity relations in the population response for the body part stimulus set. Visualization of similarity relations in the population response to the body part stimulus set as obtained using multidimensional scaling (MDS). Evoked responses (200 to 300 ms after stimulus onset) from visually responsive sites (57 sites for array P1 and 28 sites for array P2) were utilized for the MDS plot. Images containing a body part (whole bodies, heads, hands, feet) are separable from images containing an object. Additionally, images of whole bodies (blue) and heads (orange) form well defined clusters, while images of hands (red) and feet (green) are clustered together. *** P < 0.001, Pearson correlation between distances in neural space and distances in MDS space.

**Supplementary Fig S5.**
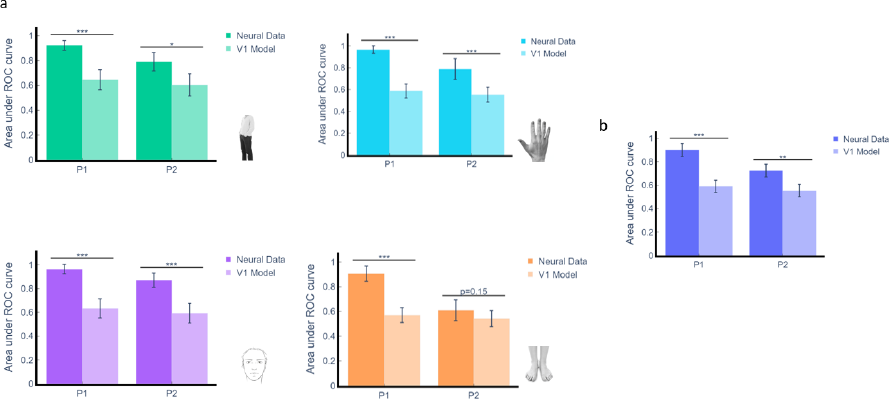
Body part selectivity control using V1 model. **a**, Comparison of performance between individual body part decoders, trained with neural data (see Fig 3b), and their corresponding site-matched models of V1-like activity. The specific body part used as the label for each decoder is indicated at the bottom right of each plot. **b**, Comparison of performance between intra-body decoders, trained with neural data (see Fig 3c), and their corresponding site-matched models of V1-like activity. Distribution for each V1 model represents 1000 iterations of randomly selecting a number of V1 units equal to the visually responsive sites of each recording session. * P < 0.05, *** P < 0.001, one-sided uncorrected permutation test.

**Supplementary Fig S6.**
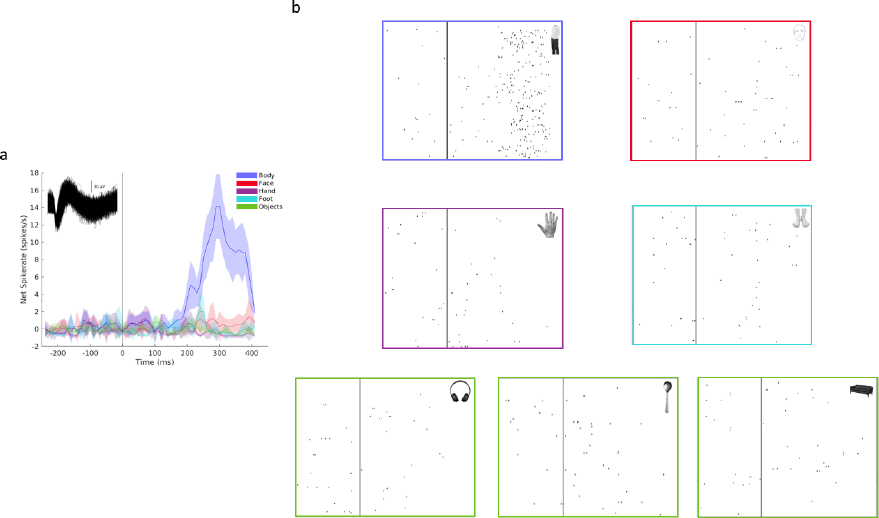
Example single unit selective for a specific body part (Specific). **a**, Time course of mean net spike rate for stimuli across the same category. Categories not containing a body part (i.e. “tools”, “manipulable objects” and “non-manipulable objects”) were grouped into a general “object” category. Vertical line corresponds to stimulus onset. The inset illustrates the spike waveform. **b,** Raster plots for individual presentations of images corresponding to each of the categories (outside rectangles colored as in (a), sample object from each category represented in top right corner).

**Supplementary Fig S7.**
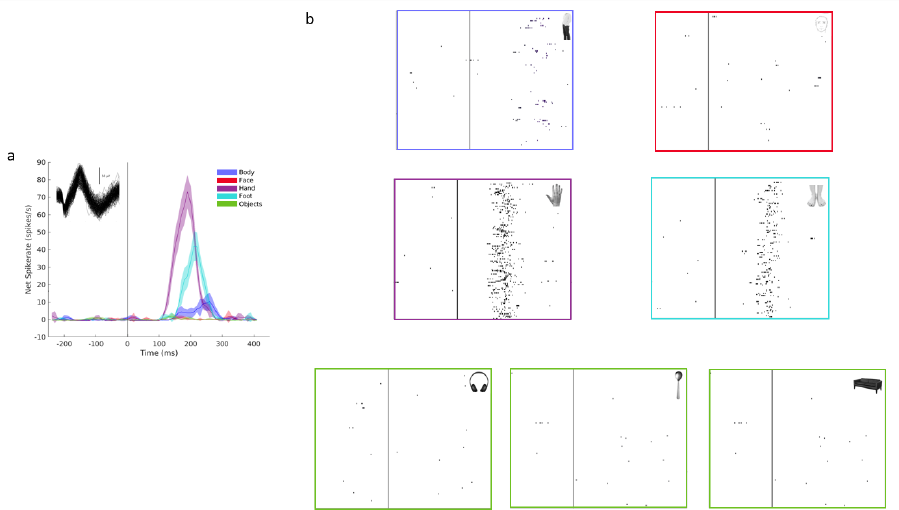
Example single unit selective for different body parts (General). **a**, Time course of mean net spike rate for stimuli across different categories. Categories not containing a body part (i.e. “tools”, “manipulable objects” and “non-manipulable objects”) were grouped into a general “object” category. Vertical line corresponds to stimulus onset. The inset illustrates the spike waveform. **b,** Raster plots for individual presentations of images corresponding to each of the independent categories (outside rectangles colored as in (a), sample object from each category represented in top right corner).

**Supplementary Fig S8.**
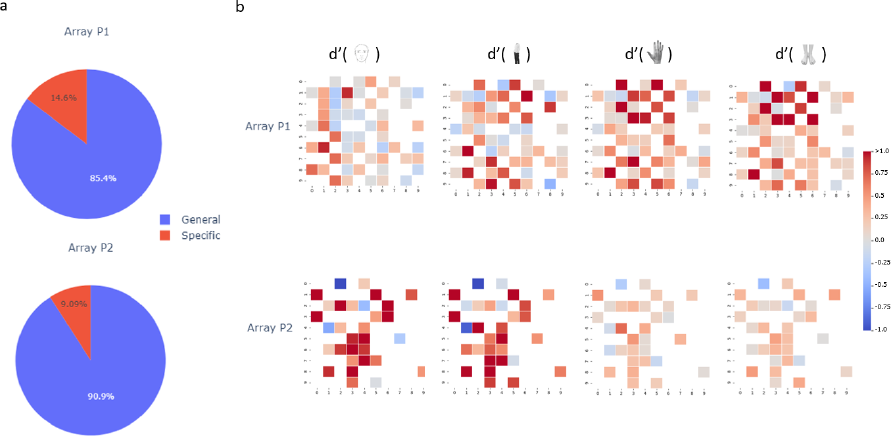
Individual site selectivity for different body parts. **a**, Distribution of population of body-selective sites with respect to the level of selectivity of each site to different body parts. Sites were categorized as either selective sites (significant d’ value for only one specific body part; example site in Supplementary Fig S6) or general sites (significant d’ value for more than one body part; example site in Supplementary Fig S7). **b,** Spatial layout of both arrays with d’ values for each body part (hands, feet, bodies without hands and feet, and heads), compared to objects.

**Supplementary Fig S9.**
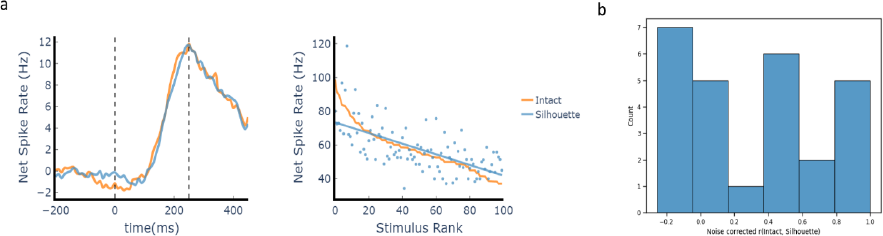
Selectivity of single sites for silhouette images. **a**, *Left panel*: Response profile of a representative site in the categorical-bodies experiment, showcasing responses to intact images (orange) and to their corresponding silhouette counterparts (blue). No significant time window was identified where the response distribution for intact images differed significantly (P < 0.01) from the silhouette versions of the images. *Right panel*: Ranked responses to intact (light orange line) and silhouette (blue dots) versions of the stimuli for the individual site in (a). Both versions were ranked based on responses to the intact stimuli. The blue line is the linear fit (ordinary least squares) to the silhouette responses based on ranking derived from the intact image responses. Spearman correlation between the distribution of responses for the intact images and their silhouette counterparts equals 0.66 with p = 5.79e-14. **b,** Distribution of noise-corrected correlations between intact and silhouette images. Each site’s noise-corrected correlation was calculated by normalizing the obtained correlation coefficient with the noise-ceiling specific to that site (Methods).

**Supplementary Fig S10.**
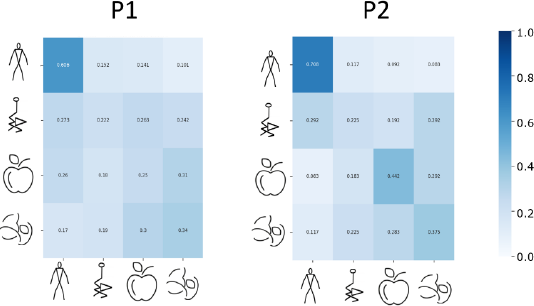
Population activity discriminates abstract bodies preferentially. Confusion matrices during the peak decoding window for the category decoding presented in Fig. 5c. The x-axis represents predicted labels, while the y-axis signifies ground-truth labels. Matrices are row-normalized to emphasize the proportion of true positives within each category. Both arrays exhibit higher discriminability for abstract bodies compared to other categories, with low misclassification rates for scrambled bodies as abstract bodies (27.3% and 29.2% for arrays P1 and P2, respectively).

**Supplementary Fig S11.**
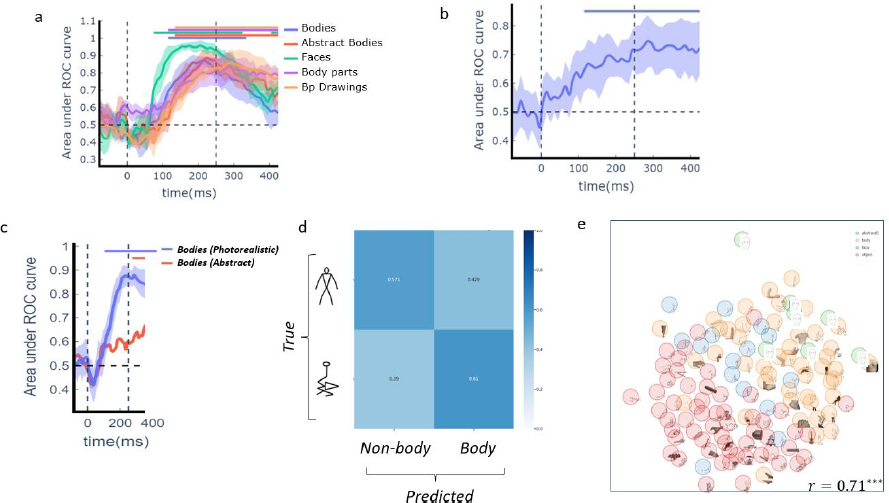
Generalization of body and abstract body representations. **a**, Comparison of time course of performance of individual decoders trained to discriminate between different body categories (labelled in top right) and objects. **b,** Time course of decoder performance for linear decoder trained to discriminate between images of abstract bodies and objects. **c,** Comparison of performance of decoders trained to discriminate between images of bodies (except faces) and objects when tested in held out data (blue) and when tested with stick figures (intact versus scrambled) (red). Lines and shading show mean ± 1std classification AU-ROC. Distribution reflects 10-fold grouped cross-validation. Horizontal bars indicate above-chance classification (P < 0.005; one-sided permutation test with at least five consecutive significant time points). **d,** Confusion matrix for generalization decoding (red line in (c)) during the peak decoding window. Predicted labels (photographic body, non-body) are on the x-axis, while ground-truth labels (stick figure body, scrambled stick figure) are on the y-axis. In this case, the data distribution used for training (photographic images) differs from that employed for generalization (stick figures). Matrices are row-normalized to emphasize the proportion of true positives within each category. **e**, Visualization of similarity relations in the population evoked responses for the reduced version of the original EBA set localizer of Downing et al. 2001 using multidimensional scaling (MDS). *** P < 0.001, Pearson correlation between distances in neural space and distances in MDS space.

**Supplementary Fig S12.**
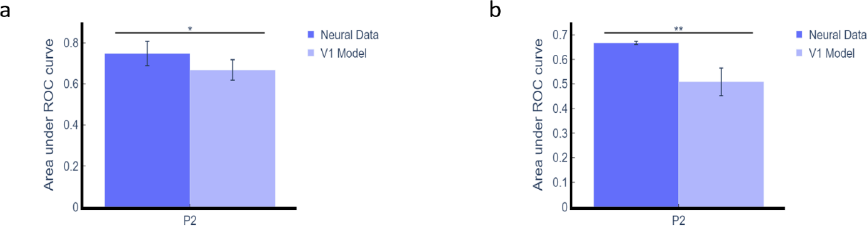
Abstract body generalization control using V1 model in array P2. **a**, Performance comparison between decoder trained with neural data to discriminate abstract bodies from non-body categories (Supplementary Fig S9b) with site-matched model employing V1-like activity. **b,** Evaluation of the generalization procedure depicted in the red line of Supplementary Fig S9c, contrasting the model trained with neural data with site-matched V1-like activity counterpart. Distribution for each V1 model represents 1000 iterations of randomly selecting a number of V1 units equal to the visually responsive sites specific to the recording session. * P < 0.05, ** P < 0.01, one-sided uncorrected permutation test.

**Supplementary Fig S13.**
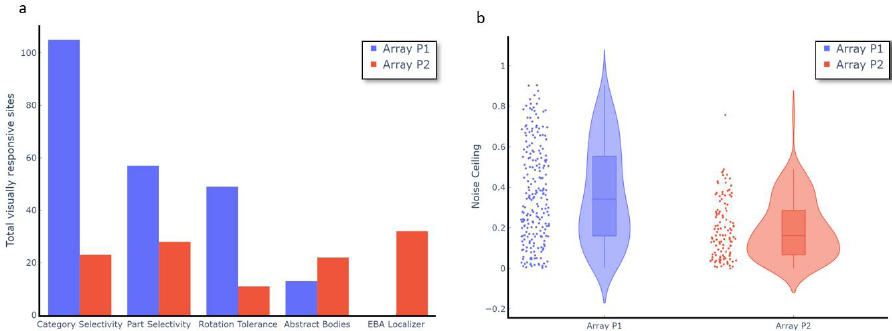
Summary of data quality across experiments. **a**, Number of visually responsive sites across sessions of the same experiment for the two arrays. **b,** Differences in the distribution of noise ceilings, pooled across all recorded sessions, for both arrays. Each individual point represents the noise ceiling of an individual visually responsive channel. The noise-ceiling of each site was computed by first computing average normalized net spike rates per image separately for each half of the data, and then measuring the (Spearman-Brown corrected) correlation between the average responses from the two halves. This entire procedure was repeated 100 times for each binary split. Violin plots represent the distribution of noise ceilings per array. Horizontal lines inside each violin plot represent the median values of each distribution (*median_*p1*_* = .34, *median_*p2*_* = .16).

## Notes

### Competing Interest Statement

The authors have declared no competing interest.

